# Computational exploration of dynamic mechanisms of steady state visual evoked potentials at the whole brain level

**DOI:** 10.1101/2021.02.05.429877

**Authors:** Ge Zhang, Yan Cui, Yangsong Zhang, Hefei Cao, Guanyu Zhou, Haifeng Shu, Dezhong Yao, Yang Xia, Ke Chen, Daqing Guo

## Abstract

Periodic visual stimulation can induce stable steady-state visual evoked potentials (SSVEPs) distributed in multiple brain regions and has potential applications in both neural engineering and cognitive neuroscience. However, the underlying dynamic mechanisms of SSVEPs at the whole-brain level are still not completely understood. Here, we addressed this issue by simulating the rich dynamics of SSVEPs with a large-scale brain model designed with constraints of neuroimaging data acquired from the human brain. By eliciting activity of the occipital areas using an external periodic stimulus, our model was capable of replicating both the spatial distributions and response features of SSVEPs that were observed in experiments. In particular, we confirmed that alpha-band (8-12 Hz) stimulation could evoke stronger SSVEP responses; this frequency sensitivity was due to nonlinear entrainment and resonance, and could be modulated by endogenous factors in the brain. Interestingly, the stimulus-evoked brain networks also exhibited significant superiority in topological properties near this frequency-sensitivity range, and stronger SSVEP responses were demonstrated to be supported by more efficient functional connectivity at the neural activity level. These findings not only provide insights into the mechanistic understanding of SSVEPs at the whole-brain level but also indicate a bright future for large-scale brain modeling in characterizing the complicated dynamics and functions of the brain.

## 1. Introduction

When our brain is stimulated by a periodic visual flickering input, a nonlinear and stimulus-locked response appears in visual processing brain regions. This neuronal response is the so-called steady-state visual evoked potential (SSVEP), and appropriately periodic visual stimuli can induce strong SSVEP responses with high signal-to-noise ratios (Pastor et al., 2003; Vialatte et al., 2010). This feature allows SSVEPs to serve as a stable paradigm to build a brain-computer interface (BCI) and estimate the characteristics of task-related neural activity (Morgan et al., 1996; Vialatte et al., 2010; Yu Zhang et al., 2013, 2015). Although SSVEPs are believed to originate in the visual cortex located in the occipital lobe, experimental studies have revealed that this evoked neural activity can be widely observed in high-level visual processing brain regions, such as the frontal and parietal lobes (Di Russo et al., 2007; Pastor et al., 2003). These findings indicate that the emergence of SSVEPs should involve multiple regions broadly distributed in the brain and that SSVEP responses might thus be modulated by fundamental properties of brain networks. Additionally, SSVEPs have also been found to exhibit strong frequency sensitivity; moreover, in particular, they respond optimally to a suprathreshold stimulus at the alpha frequency band (8-12 Hz) (Norcia et al., 2015; Xu et al., 2013). However, despite accumulating experimental data, the biophysical mechanisms of both the frequency sensitivity of SSVEPs and the regulation of SSVEP responses in the brain remain largely unexplored.

Recent studies using computational modeling have provided deep insights into the mechanistic understanding of complicated brain dynamics and functions (Gosak et al., 2018; Parastesh et al., 2021). In this filed, most of theoretical investigations on SSVEPs tried to reproduce nonlinear SSVEP dynamics at the neural circuit level (Labecki et al., 2016; Yang et al., 2019). Using a neural-field model of the cortex and thalamus, Roberts and Robinson showed the fundamental spectral properties of our brain when prompted by periodic visual stimuli (Roberts & Robinson, 2012). This physiologically based model not only explains the entrainment and harmonic behaviors of SSVEP responses but also predicts rich nonlinear dynamics in response to stimuli with high suprathreshold amplitudes. Further investigation revealed that several key features of SSVEP spectra can be captured by a simplified neural mass model consisting of excitatory and inhibitory neural populations. Using such an ideal model, it has been indicated that the harmonic and subharmonic components of SSVEPs are a natural consequence of the nonlinearities of neural populations (Labecki et al., 2016). Moreover, recent computational studies also suggested that the response of SSVEPs can be modulated by the interactions between different brain regions. By constructing a laminar cortical model that is composed of the primary visual cortex (V1) and the secondary visual cortex (V2), we have shown that SSVEP modulation is implemented by alpha oscillation in a complementary manner at different spatial levels (Yang et al., 2019). In particular, it is found that interlaminar coupling contributes to the laminar-specific organization of the evoked response following the opposite rules in the intracortical and intercortical drive (Yang et al., 2019), which unifies experimental observations that originally seemed contradictory (Koch et al., 2008; Morgan et al., 1996). To our knowledge, however, these modeling studies mainly focused on local neural circuits and did not consider exploring large-scale brain dynamics of SSVEPs using computational models under realistic connectivity constraints.

Remarkably, the rapid development of large-scale brain modeling offers a powerful approach to reveal the neural mechanisms of specific cognitive functions or stimulus-induced activity at the whole-brain level (Breakspear, 2017; Gosak et al., 2018; Ponce-Alvarez et al., 2015; Shine et al., 2018). As pioneering work, a model of the large-scale macaque cortex was developed to explore spatiotemporal features of spontaneous cortical dynamics with anatomical constraints (Honey et al., 2007, 2009). During the resting state, this model exhibited several rich and interrelated spatiotemporal structures at multiple time scales, thus indicating that functional connectivity (FC) may be significantly shaped by brain structure. To further promote model performance, theoretical researchers have proposed constructing large-scale brain models by combining both the FC and structural connectivity (SC) acquired using magnetic resonance imaging (MRI) techniques (Gustavo Deco et al., 2019; Demirtaş et al., 2018). In particular, functional MRI (fMRI) data have been widely considered a condition of constrained optimization during model establishment. Such inverse-based models are capable of reproducing important features of large-scale brain dynamics, mainly because they link the structural and functional organization of the brain together (Cabral et al., 2017; G. Deco et al., 2014). Importantly, it has also been shown that the dynamic features of these large-scale brain models are governed by parameters with appropriate biophysical interpretation (Joglekar et al., 2018; Schirner et al., 2018). This fact allows several critical dynamical behaviors generated by large-scale models to be verified in well-designed experiments. Therefore, such large-scale brain modeling techniques have unique advantages that bridge system-level neural dynamics and specific cognitive functions or stimulus-induced activity in the brain.

In this study, we investigated the dynamic mechanisms of SSVEPs by constructing a large-scale brain model with human SC and FC data. To this end, the model was optimized with an iterative-fitting strategy proposed by Deco et al (G. Deco et al., 2014). By stimulating the model with periodic visual input in early-stage visual processing brain regions (i.e., the occipital lobe), we found that the stimulus-evoked potential could be propagated within our optimized model and SSVEP responses could also be detected in high-level visual processing areas, such as several regions in the frontal lobe. In particular, SSVEPs were optimal in response to a suprathreshold stimulus in the alpha band, and such frequency sensitivity of SSVEPs was thought to be a consequence of entrainment and resonance (Herrmann et al., 2016; Lab Notbohm et al., 2016). Additionally, the performance of SSVEP responses was also significantly related to the network properties, and a better SSVEP performance corresponded to a higher stimulus-evoked FC efficiency at the neural activity level. Our results thus provide a new perspective to understand the nonlinear responses of SSVEPs within the whole brain framework.

## 2. Model and Methods

### 2.1 Empirical structural and functional connectivity

A standard MRI dataset consisting of both empirical structural connectivity (SC) and functional connectivity (FC) was employed to establish the large-scale brain model (G. Deco et al., 2014; Hagmann et al., 2008). Briefly, the SC and FC data in this MRI dataset were derived from five healthy right-handed male human participants (age 29.4 ± 3.4 years) and were acquired with a Philips Achieva 3T MRI system. For each subject, diffusion spectrum imaging (DSI) was performed to track white matter tracts; then, the empirical SC matrix was constructed with the anatomical landmarks of 66 gray matter cortical regions, which are listed in Table 1 (G. Deco et al., 2014; Hagmann et al., 2008). Theoretically, each SC element represents the connectivity density between a pair of cortical regions. The average empirical SC matrix among all subjects was the connectivity matrix used to couple different brain regions. However, we set the connection of a region to itself to 0 in the connectivity matrix because the effect of internal interactions was already considered in the microcircuit structure for each brain region (see below).

**Table 1.** Names and abbreviations for the 66 cortical regions used in the present study. Two labels (i.e., R and L) refer to the right and left hemispheres, respectively.

| Region name | Abbreviation | Label (R) | Label (L) |
| --- | --- | --- | --- |
| Bank of the superior temporal sulcus | BSTS | 12 | 55 |
| Caudal anterior cingulate cortex | CAC | 23 | 44 |
| Caudal middle frontal cortex | CMF | 17 | 50 |
| Cuneus | CUN | 29 | 38 |
| Entorhinal cortex | ENT | 1 | 66 |
| Frontal pole | FP | 4 | 63 |
| Fusiform gyrus | FUS | 5 | 62 |
| Inferior parietal cortex | IP | 10 | 57 |
| Inferior temporal cortex | IT | 9 | 58 |
| Isthmus of the cingulate cortex | ISTC | 31 | 36 |
| Lateral occipital cortex | LOCC | 7 | 60 |
| Lateral orbitofrontal cortex | LOF | 22 | 45 |
| Lingual gyrus | LING | 27 | 40 |
| Medial orbitofrontal cortex | MOF | 26 | 41 |
| Middle temporal cortex | MT | 13 | 54 |
| Paracentral lobule | PARC | 30 | 37 |
| Parahippocampal cortex | PARH | 2 | 65 |
| Pars opercularis | POPE | 18 | 49 |
| Pars orbitalis | PORB | 21 | 46 |
| Pars triangularis | PTRI | 19 | 48 |
| Pericalcarine cortex | PCAL | 28 | 37 |
| Postcentral gyrus | PSTS | 15 | 52 |
| Posterior cingulate cortex | PC | 33 | 34 |
| Precentral gyrus | PREC | 16 | 51 |
| Precuneus | PCUN | 32 | 35 |
| Rostral anterior cingulate cortex | RAC | 24 | 43 |
| Rostral middle frontal cortex | RMF | 20 | 47 |
| Superior frontal cortex | SF | 25 | 42 |
| Superior parietal cortex | SP | 8 | 59 |
| Superior temporal cortex | ST | 14 | 53 |
| Supramarginal gyrus | SMAR | 11 | 56 |
| Temporal pole | TP | 3 | 64 |
| Transverse temporal cortex | TT | 6 | 61 |

We used the empirical FC matrix as the gold standard to optimize the large-scale brain model. For each subject, blood oxygenation level-dependent (BOLD) signals (20 mins) were obtained in the resting-state using the same 66 cortical regions described above (Buxton & Frank, 1997). A global mean signal was regressed out from the BOLD signals before the FC was calculated. Then, the resting-state FC matrix was constructed for each subject by measuring the Pearson’s correlation of BOLD signals among different brain regions. We averaged the resting-state FC matrix across all subjects as the final empirical FC matrix in the present study (for details; see Honey et al., 2009).

### 2.2 Large-scale computational model of the brain

To explore the dynamic mechanisms of SSVEPs at the whole-brain level, we established a large-scale model of the brain composed of 66 cortical regions. Regions in the model were initially coupled by the empirical SC matrix. As schematically shown in Fig. 1A, the microcircuit of each brain region was assumed to be a local network of excitatory and inhibitory populations, and their dynamic behaviors were characterized by Wilson-Cowan equations. The dynamics of our large-scale brain model are described as follows:

**Figure 1.**
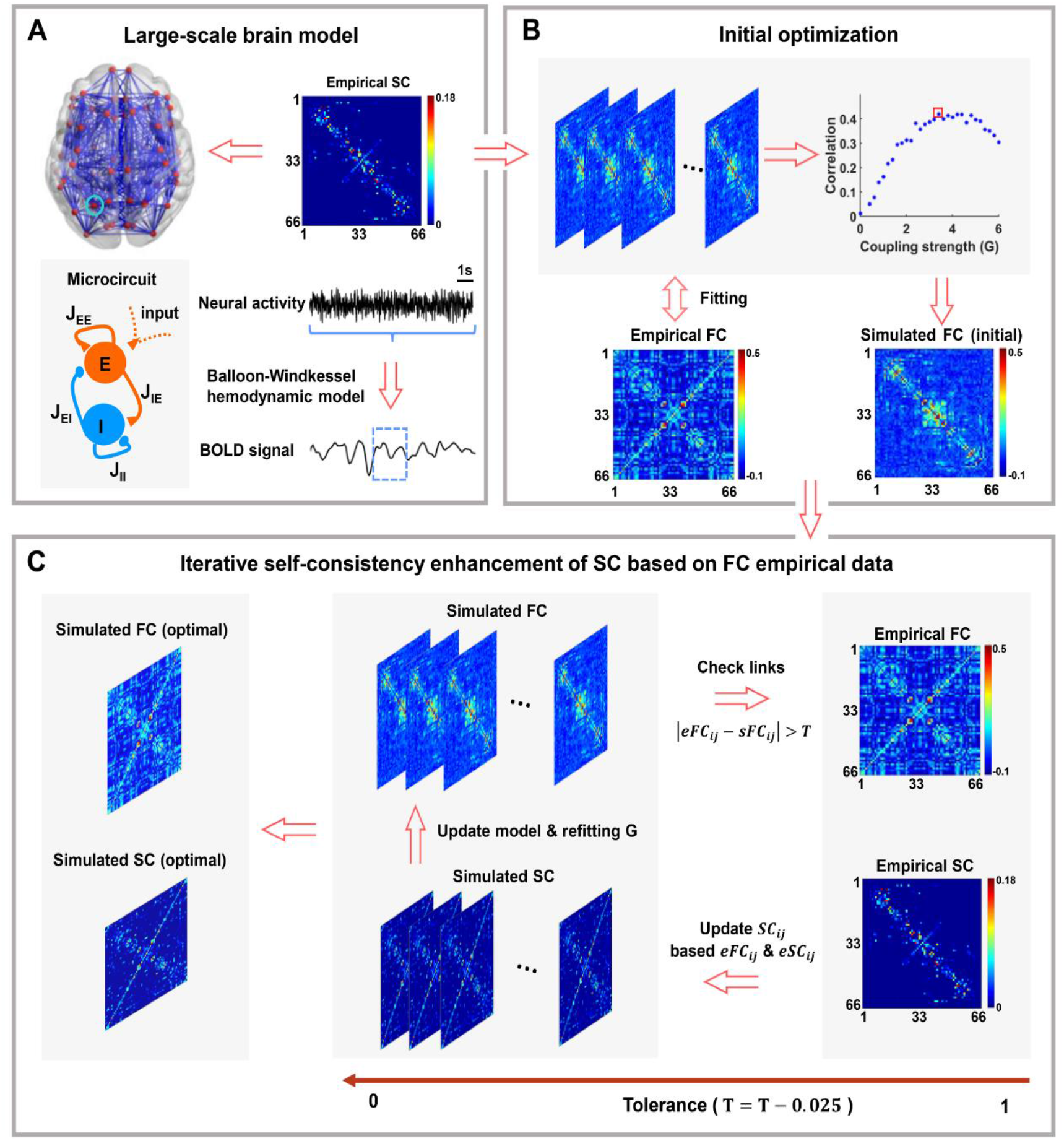
Overview of the large-scale brain model. **A:** In the large-scale brain model, each brain region is modeled as a microcircuit that is composed of coupled excitatory (E) and inhibitory (I) neural populations. The empirical SC is used to define initial connectivity among different brain regions. The excitatory neural activity can be converted into the simulated BOLD signal with the Balloon-Windkessel hemodynamic model. As an example, the blue dotted square represents the BOLD signal transformed from the above neural activity. **B:** Initial optimization for the large-scale brain model. After fitting, an initial simulated FC is obtained at an optimal global coupling factor *G* (red square) under the constraints of empirical SC and FC. **C:** Schematic presentation of iterative self-consistency enhancement of SC based on empirical FC. This strategy begins with an initial tolerance of 1 and stops when the tolerance level is close to 0. A detailed description of this optimization strategy can be found in the Model and Methods section. With the iterative-fitting strategy, we obtained both the optimal simulated SC and FC used in this study.

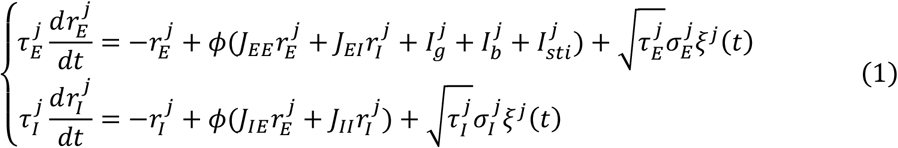

where *j* indexes different brain regions, 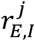 represents the mean firing rate of the excitatory (E) and inhibitory (I) populations of the *j* -th region, 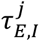 denotes the corresponding time constants, and *ξ*^*j*^(*t*) is Gaussian white noise with zero mean and standard deviation 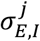 for excitatory and inhibitory neural populations in the *j* -th region. The transduction function *ϕ*(*x*) = *x*/(1 − *e*^−*x*^) is employed to convert the current *x* to the firing rate. In our model, the synaptic inputs within the microcircuit (i.e., 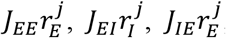and 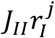) are governed by four synaptic coupling variables *J*_*EE*_, *J*_*EI*_, *J*_*IE*_, and *J*_*II*_. A background input 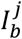 was fed to each excitatory population to maintain spontaneous brain activity. Each excitatory population also received the global synaptic input from other brain regions according to the function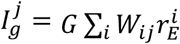, where the outer sum runs over interconnections onto the particular region *j, W*_*ij*_ is an element of the SC matrix representing the coupling between the regions *i* and *j*, an *G* is a global coupling factor (also termed as global-scale coupling) that requires optimization. In addition, excitatory populations in several specific visual areas were driven by an external stimulus 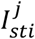 to induce a stimulus-evoked brain state.

Note that our model was first optimized with an iterative-fitting strategy (see below) in the resting state, and then the optimized large-scale brain model was used to investigate the dynamic mechanisms of SSVEPs in the stimulus-evoked state. When the brain received a uniform flash stimulus, SSVEPs could be recorded in a variety of visual areas (Rager & Singer, 1998). To excite SSVEP responses in the model, we injected periodic visual input as an external stimulus into the occipital-related regions, including the lateral occipital cortex (LOCC), pericalcarine cortex (PCAL), lingual gyrus (LING) and cuneus (CUN). In simulations, this external periodic stimulus was modeled as the square wave described by:

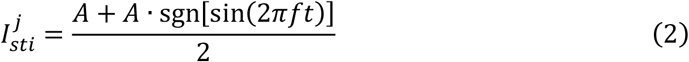

Here, *A* and *f* represent the amplitude and frequency of the stimulus, and sgn(·) is the sign function. Additionally, we have demonstrated the similar SSVEP responses can be also elicited by other types of external periodic stimuli, such as the flicking input with a sinusoidal wave profile (Supplementary Fig.1).

In simulations, stochastic differential equations were integrated by using the Euler-Maruyama method with a time step of 0.1 ms. Unless mentioned otherwise, the default values of the model parameters chosen are those presented in Table 2.

**Table 2.** The default values of the model parameters used in the simulations.

| Parameters | Description | Values |
| --- | --- | --- |
| $\tau_E^j$ | Excitatory synaptic time constants | 18 ms |
| $\tau_I^j$ | Inhibitory synaptic time constants | 25 ms |
| $I_b^j$ | Background current | 2 |
| $A$ | Amplitude of the stimulus signal | 0.5 |
| $\sigma_E^j$ | Noise strength of the excitatory population | 0.45 |
| $\sigma_I^j$ | Noise strength of the inhibitory population | 0.45 |
| $J_{EE}$ | Synaptic strength of $E \rightarrow E$ coupling | 1.5 |
| $J_{EI}$ | Synaptic strength of $I \rightarrow E$ coupling | -2.6 |
| $J_{IE}$ | Synaptic strength of $E \rightarrow I$ coupling | 3.5 |
| $J_{II}$ | Synaptic strength of $I \rightarrow I$ coupling | -2.5 |

### 2.3 Model optimization with an iterative-fitting strategy

We employed an iterative-fitting strategy to optimize the large-scale brain model in the resting state (G. Deco et al., 2014). This optimal strategy is based on enhancing the original SC matrix by adding new links for pairs of nodes according to the corresponding FC between those nodes (G. Deco et al., 2014). Before the iterative-fitting process, we first constructed the large-scale brain model with the empirical SC matrix (Fig. 1B) and searched for an optimal global coupling factor *G* by maximizing the correlation between the empirical FC (Fig. 1B) and the simulated FC. To compute the simulated FC, we ran each simulation for 500 s to generate data for a sufficient period of time at the neural activity level and removed the first 20 s of data before analysis. Then, the Balloon-Windkessel hemodynamic model (Supplementary Text 1) was used to convert the neural activity of the excitatory population into the simulated BOLD signal (Buxton & Frank, 1997; Buxton et al., 1998; Friston et al., 2003). The simulated BOLD signal was downsampled to a low frequency (2 Hz) to have temporal resolution comparable with that of the empirically measured fMRI recordings. For each simulation, the simulated FC was obtained by computing the Pearson’s correlation of the simulated BOLD signals among different brain regions. We averaged the simulated FC over 5 trials as the final simulated FC. An initial optimization is performed for our model by varying the global coupling factor *G*, and the optimal strength of *G* is identified by maximizing the correlation between the empirical FC and the simulated FC (Fig. 1B). After this initial optimization, an initial simulated FC could be determined at an optimal strength of the global-scale coupling *G* under constraints of empirical structural and functional connectivity (Fig. 1B).

With the iterative-fitting strategy, we tried to further improve the similarity between the empirical and simulated FC by adding a few links to the empirical SC. To do this, the maximal value of the empirical FC matrix was normalized to 1, and an initial tolerance level was set to 1 to control the iterative process. We started the iterative-fitting algorithm with conditions under the normalized empirical FC as well as an initial simulated FC based on the empirical SC at the optimal strength of *G* (Fig. 1C). The iteration process can be mathematically described as follows: we defined the matrix of the simulated FC and normalized empirical FC as *sFC* and *eFC*, respectively. At each iteration step, we needed to identify all connections in the simulated FC and normalized empirical FC satisfying the judgement condition of |*eFC*_*ij*_ − *sFC*_*ij*_| > *T*. When *eFC*_*ij*_ > 0, the SC links for those identified connections were updated with the following rule: *SC*_*ij*_ = 0.15 · *eFC*_*ij*_. In the case of *eFC*_*ij*_ ≤ 0, the corresponding SC links were redefined as *SC*_*ij*_ = 0 when it was zero in the original matrix; otherwise, *SC*_*ij*_ was scaled down to a minimum value of weights (0.0005) as described in the original matrix (G. Deco et al., 2014). Then, the large-scale brain model was reoptimized by maximizing the correlation between the empirical FC and the simulated FC with the newly generated SC versus the global coupling factor *G*. The above process was iterated several times by reducing the tolerance level (*T* = *T* − 0.025) and was stopped when the tolerance level equals to 0.025. It is worth noting that the stopped tolerance level should be larger than 0 due to the condition of |*eFC*_*ij*_ − *sFC*_*ij*_| > *T*. Using this iterative-fitting strategy, the maximum fit between the optimal simulated SC and FC could be identified at an appropriate tolerance level (Fig. 1C). Under this condition, the correlation between the empirical and simulated FC achieved its maximal value, and the corresponding tolerance level is a judgement criterion for our model optimization. Compared with intrahemispheric connectivity, the anatomical structure of the brain derived from the DSI data would miss a relatively larger number of long-range interhemispheric connections. As reported previously (G. Deco et al., 2014), the iterative self-consistency enhancement of SC based on empirical fMRI data mainly increases the interhemispheric connections between homologous areas and can significantly promote the performance of the large-scale brain model.

### 2.4 Data analysis

We used several measurements to quantify SSVEP performance. Since SSVEPs are a fast stimulus-evoked response in the brain, high-temporal resolution data are required for capturing the rapid dynamics of SSVEPs. Therefore, in addition to model optimization, all other data generated by our model were analyzed at the neural activity level. For each experimental condition, we ran simulations of 20 trials with different random seeds. For each trial, the simulation was carried out for 500 s with stochastic initial conditions, and the first 20 s of data were removed before analysis. The rest of the data were resampled to 250 Hz, which is comparable to the sampling rate of real electroencephalography (EEG) recordings. These 480-s-long data epochs were further divided into 48 segments, with each segment lasting for 10 s. In the following studies, we used all 48 data segments recorded to analyze SSVEP responses and randomly chose 5 data segments when measuring network properties and network synchronization. For each experimental condition, we calculated the segment-averaged results for each trial and then reported the data across 20 trials as the final results.

#### 2.4.1 Analysis of SSVEP responses

To measure the performance of SSVEP responses to the periodic driven stimulus, both the power and signal-to-noise ratios (SNRs) were estimated with the power spectral analysis for specific areas within the occipital and frontal lobes. These brain areas are highly associated with early and late stages of visual processing, and the SSVEPs have been widely observed in these regions. In this analysis, we computed the power spectrum density of neural activity in these visual-related regions by using the fast Fourier transform (FFT) method (Srinivasan et al., 2006). The SSVEP power *S*(*f*_0_) was simply defined as the corresponding amplitude of the power spectral density at the stimulus frequency *f*_0_. Then, the SNR value of the stimulus-evoked response was calculated as follows:

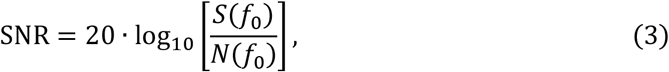

where *f*_0_ is the stimulus frequency and 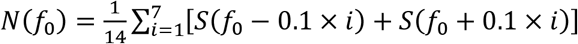 represent the average power near the stimulus frequency *f*_0_ (1.4 Hz band centered on the stimulus frequency but excluding the stimulus frequency itself; the frequency resolution is 0.1 Hz). Additionally, to visualize the features of neural activity in both the time and frequency domains, time-frequency analysis was performed with the wavelet method for several areas located in the occipital and frontal lobes. The classical Morlet wave was used as the wavelet basis function, and the default bandwidth parameter and wavelet center frequency were fixed at 0.5 Hz and 1 Hz, respectively.

#### 2.4.2 Analysis of network properties

To quantify the SSVEP performance at the network level, we constructed the weighted brain networks under the resting state and stimulus-evoked state. For both types of brain states, we measured the FC among different regions at the neural activity level with coherence, which was defined as follows (Nunez et al., 1997):

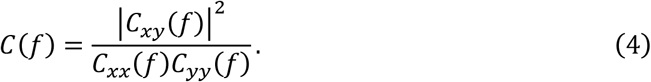

Here *C*_*xy*_(*f*) is the cross-spectrum between the neural activity of excitatory populations *x*(*t*) and *y*(*t*) from different brain regions, and *C*_*xx*_(*f*) and *C*_*yy*_(*f*) are the corresponding autospectra at frequency *f*.

Several measurements were employed to assess network properties under different brain states. These network properties have been widely used in previous studies on brain network analysis and include the clustering coefficient, characteristic path length, global efficiency, and local efficiency (Newman, 2003; Watts & Strogatz, 1998). In this study, we used the Brain Connectivity Toolbox (www.brain-connectivity-toolbox.net) to calculate these network properties, with their detailed mathematical descriptions provided below.

To evaluate the degree of network collectivization, we calculated the clustering coefficient of the weighted brain network as follows:

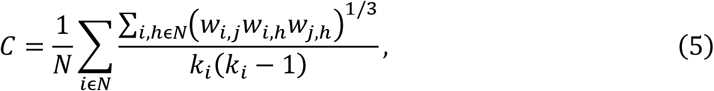

where *N* is the number of nodes in the network, *w*_*i,j*_ indicates the weight between nodes *i* and *j*, and *k*_*i*_ is the degree of node *i*.

The characteristic path length *L* is defined as the average of the shortest path length *L*_*ij*_ between any two nodes in the network. Mathematically, this measurement can be computed as:

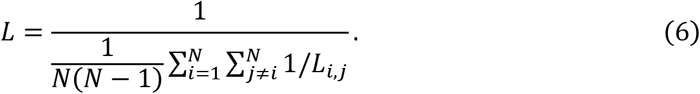

Moreover, we estimated both the global and local efficiency of brain networks under the resting state and stimulus-evoked state. Global efficiency is defined as (Latora & Marchiori, 2001):

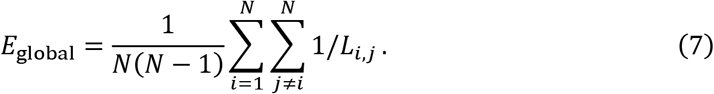

Theoretically, a smaller shortest path length (or a larger clustering coefficient) corresponds to a higher global efficiency with a relatively faster information transfer between nodes in the network. Compared with global efficiency, local efficiency reflects the extent of integration between the immediate neighbors of the given node (Achard & Bullmore, 2007). By definition, local efficiency could be obtained by averaging the local efficiencies of all nodes in a network *G*.

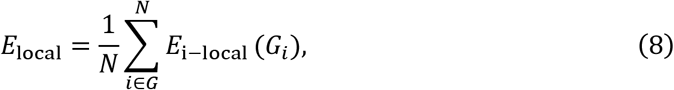

with

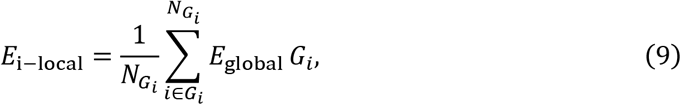

where *G*_*i*_ denotes the subgraph comprising all nodes that are immediate neighbors of node *i*, and 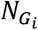 is the number of nodes in *G*_*i*_.

#### 2.4.3 Measurement of network synchronization

The synchronization of neural activity among different brain regions is believed to play a crucial role in highly efficient neuronal information and cognitive processing (Della Rossa et al., 2020; Melloni et al., 2007; Parastesh et al., 2021). In this study, we also compared the synchronization of neural activity generated by our large-scale brain model under different brain states. Based on the mean-field theory, the synchronization factor *R* for the network can be mathematically calculated as:

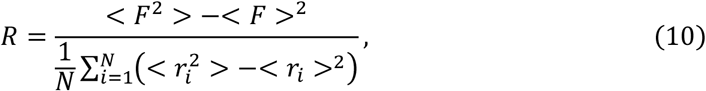

where *r*_*i*_ is the mean firing rate of the excitatory population of the *i*-th region, *N* is the total number of brain regions, 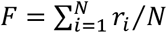 is the average neural activity across all brain regions, and the symbol <·> represents the mean of the variable over time. Theoretically, the synchronization factor *R* is within the range [0, 1], and a larger *R* indicates a relatively higher level of synchronous neural activity among different brain regions.

## 3. Results

### 3.1 Optimization of the large-scale brain model

In this study, we investigated the dynamic mechanisms of SSVEPs using a large-scale brain model. As a preliminary step, we optimized the model with constraints of realistic human imaging data, and allowed it to generate simulated BOLD signals that can be comparable with real fMRI recordings. For this purpose, initial optimization was performed for the model by using the empirical SC and FC (Fig. 1B). At an optimal global-scale coupling of *G* = 3.41, simulated FC derived from our large-scale brain model showed the best match with empirical FC (red square in Fig. 1B). Under this condition, the correlation between simulated and empirical FC achieved the maximal value of 0.41. To further improve the fit between simulated and empirical FC, we introduced an iterative-fitting strategy proposed in a previous study (G. Deco et al., 2014), which is systematically summarized in Fig. 1C. At the initial tolerance level of *T* = 1, we started this iterative-fitting strategy with the best fitted FC corresponding to the original empirical SC. During the iteration process, additional links were added into the SC matrix to reduce the tolerance level, and the new simulated SC was updated at each step. At a relatively low level of tolerance (*T* = 0.125), the large-scale brain model exhibited the best performance at an optimal simulated SC (Fig. 2A). To achieve this optimal fit, we observed that the optimal global-scale coupling of *G* was fixed at 3.01. By comparing simulated SC with empirical SC, we identified approximately 10.28% and 21.25% newly added intrahemispheric and interhemispheric connections within the optimal simulated SC matrix (Figs. 2B and 2C). To our surprise, the correlation between the optimal simulated and empirical FC was improved to 0.82 (Figs. 2A and 2C). Our results further demonstrate the superiority of the iterative-fitting strategy, showing that the optimized model can best reproduce the large-scale brain dynamics by adding a certain number of new links to the SC derived from the DSI data.

**Figure 2.**
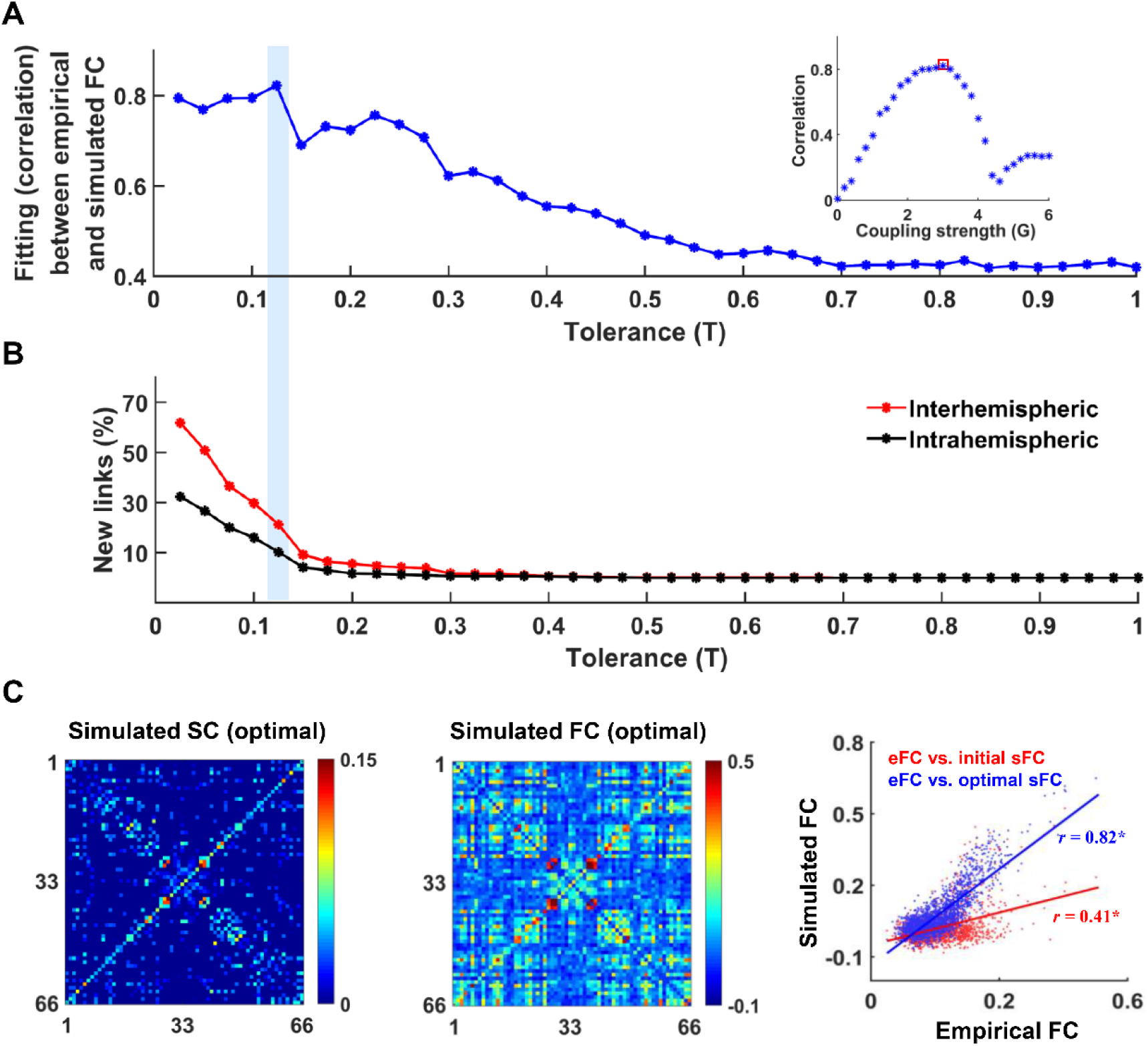
Model optimization based on the iterative-fitting strategy. **A:** The fitting between empirical and simulated FC with decreasing tolerance *T*. The optimal simulated FC is obtained at an optimal global coupling factor *G* (red square) under the constraints of the new simulated SC (updated SC at tolerance *T* = 0.125). **B:** The cumulative percentage of new intrahemispheric (black) and interhemispheric (red) links added to simulated SC with decreasing tolerance. **C:** The optimal simulated SC (left panel) and FC (middle panel) matrices are obtained at a relatively low level of tolerance (*T* = 0.125) and an optimal global-scale coupling factor (*G* = 3.01). The right panel shows correlations between empirical and simulated FC before and after the iterative-fitting strategy. The red line indicates the correlation between empirical FC and initial simulated FC (*r* = 0.41), and the blue line shows the correlation between empirical FC and optimal simulated FC obtained by using an iterative-fitting strategy (*r* = 0.82). *r* denotes the correlation coefficient, and the symbol * means significant at the 99% level (*p* < 0.01). Statistical significance was determined by the two-tailed Student’s t-test.

### 3.2 Typical SSVEP responses can be elicited in a large-scale brain model

To examine whether the optimized large-scale brain model can capture the fundamental features of SSVEP responses, we excited the occipital lobe (LOCC, PCAL, LING, and CUN) with an external periodic stimulus and detected SSVEP responses in both these occipital regions and several frontal-related regions (frontal pole (FP), pars orbitalis (PORB), lateral orbitofrontal cortex (LOF), and medial orbitofrontal cortex (MOF)) (Fig. 3A and Table 1). From a functional perspective, the occipital lobe is involved primarily in the early stage of visual processing, whereas the abovementioned frontal regions are believed to participate in higher visual processing. In this work, the external periodic stimulus was generated by a square wave with default parameter values (amplitude 0.5 nA and frequency 10 Hz).

**Figure 3.**
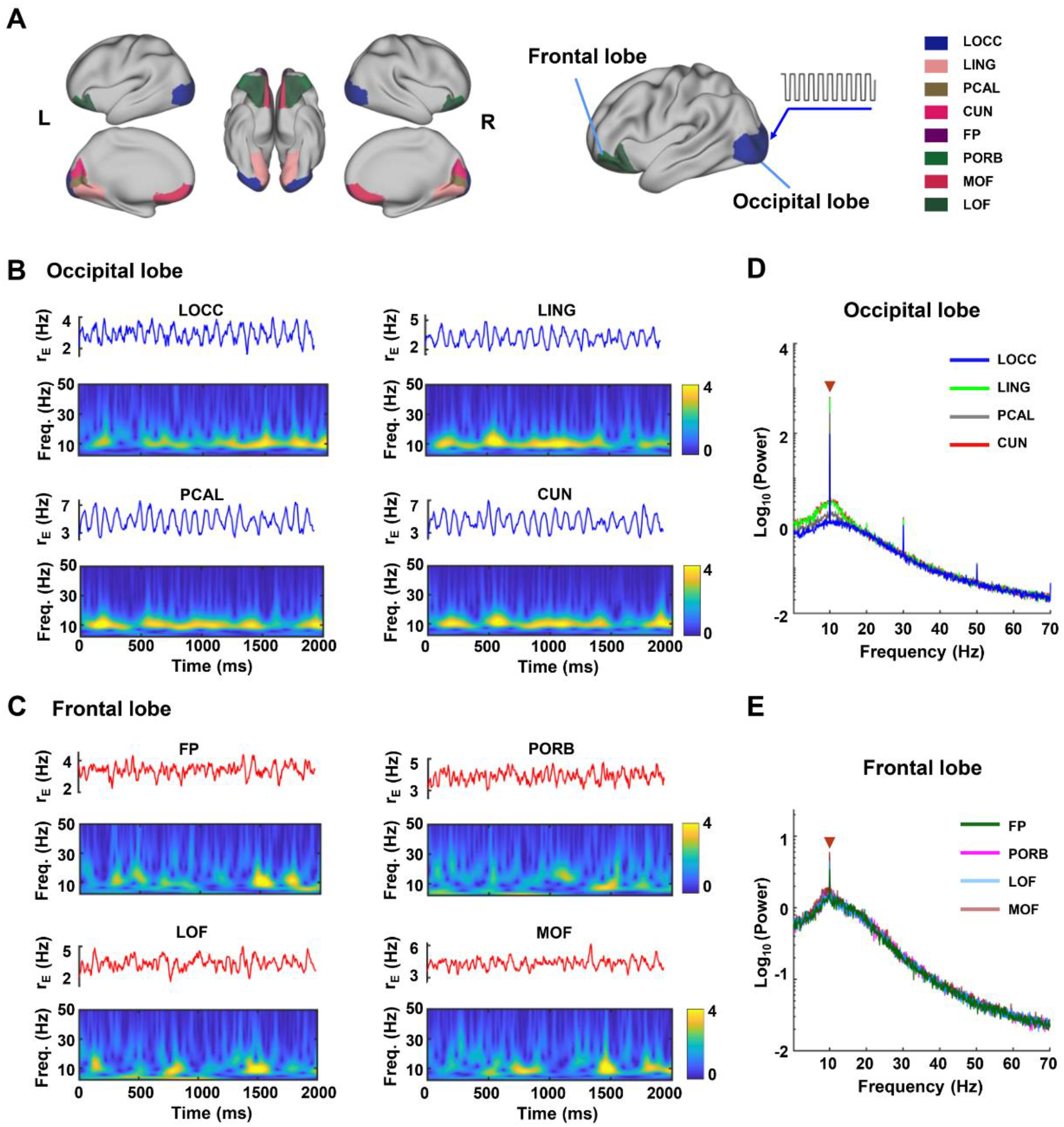
SSVEP responses at different occipital and frontal regions. **A:** Schematic diagram of the occipital-related regions (LOCC, LING, PCAL and CUN) and frontal-related regions (FP, PORB, LOF and MOF) evaluated in this study. To simulate flickering visual stimulation, these occipital regions were assumed to be driven by a square wave with an amplitude of 0.5 nA and a frequency of 10 Hz. **B, C:** Examples of typical neural activity and the corresponding time-frequency spectrogram for brain regions distributed in the occipital (B) and frontal lobes (C). **D, E:** The average power spectrum of neural activity for regions in the occipital (D) and frontal (E) lobes. The red triangle denotes the SSVEP peaks occurring at a stimulus frequency of 10 Hz. The large-scale brain model can reproduce the fundamental dynamic features of SSVEP responses.

In Figs. 3B and 3C, we show typical examples of neural activity and the corresponding time-frequency spectrogram for different occipital and frontal regions, respectively. For each time-frequency spectrogram, a remarkable power increase could be observed near the driving frequency of the external periodic stimulus. This indicated that SSVEPs could be elicited in both the occipital and frontal regions, two main sources of SSVEPs observed in experimental studies. Consistent with experimental studies, brain regions distributed in the occipital lobe showed much stronger SSVEP responses than frontal-related regions (Ding et al., 2006; Morgan et al., 1996; Srinivasan et al., 2007). This is not surprising because SSVEPs are believed to be originally elicited in the occipital area, and its propagation through nerve fibers may lead to a notable reduction in evoked power. Further power spectrum analysis with FFT not only revealed an obvious power peak located at 10 Hz for each region but also demonstrated distinct SSVEP performance in the occipital-related regions (Figs. 3D and 3E). Compared with other occipital regions, the LOCC was a mid-level visual processing area and displayed relatively lower SSVEP power (Fig. 3D). In addition, more complicated features of the SSVEP spectra, including odd harmonic components, could also be observed in these occipital regions (Fig. 3D). Theoretically, this might be because the square wave input contains only odd harmonics. However, due to signal attenuation during the propagation process, such complicated spectral features of SSVEP responses disappeared in the frontal lobe (Fig. 3E). Overall, the above results confirmed that the large-scale brain model can reproduce the fundamental dynamic features of SSVEP responses.

### 3.3 Impacts of an external periodic stimulus on SSVEP responses

Previous experimental studies have documented that the performance of SSVEP responses could be greatly impacted by the physical properties of periodic visual stimuli (Ding et al., 2006; Morgan et al., 1996). As important nonlinear SSVEP dynamics, it has been specifically reported that responses of SSVEPs in the occipital and frontal cortex are strongly sensitive to the frequency of visual stimuli (Di Russo et al., 2007; Labecki et al., 2016; Srinivasan et al., 2006). Using our large-scale brain model, we studied the dependence of SSVEP performance on the stimulus frequency of the external periodic visual input. In Figs. 4A and 4B, we depicted both the power and SNR of SSVEPs as a function of the stimulus frequency for different occipital regions. With increasing stimulus frequency, we found that both the SSVEP power and the SNR value first rose and then dropped. For each brain region, their maximal values were achieved at an intermediate stimulus frequency. In agreement with experimental observations (Ding et al., 2006; Xu et al., 2013), our large-scale brain model correctly predicted that these occipital regions would show optimal responses to external periodic stimuli in the alpha frequency band (8-12 Hz). This nonlinear SSVEP behavior is the so-called frequency sensitivity, and similar findings were also observed in the frontal regions (Figs. 4C and 4D). However, in comparison with the occipital lobe, both SSVEP power and SNR values observed in these frontal regions showed significantly lower magnitudes at different stimulus frequencies (Mann–Whitney U test, *p* < 0.001). In addition, the frequency-sensitivity range was slightly reduced during the propagation of evoked neural activity. Such a reduction resulted in a relatively narrow frequency-sensitivity range for regions distributed in the frontal lobe (Fig. 4D).

**Figure 4.**
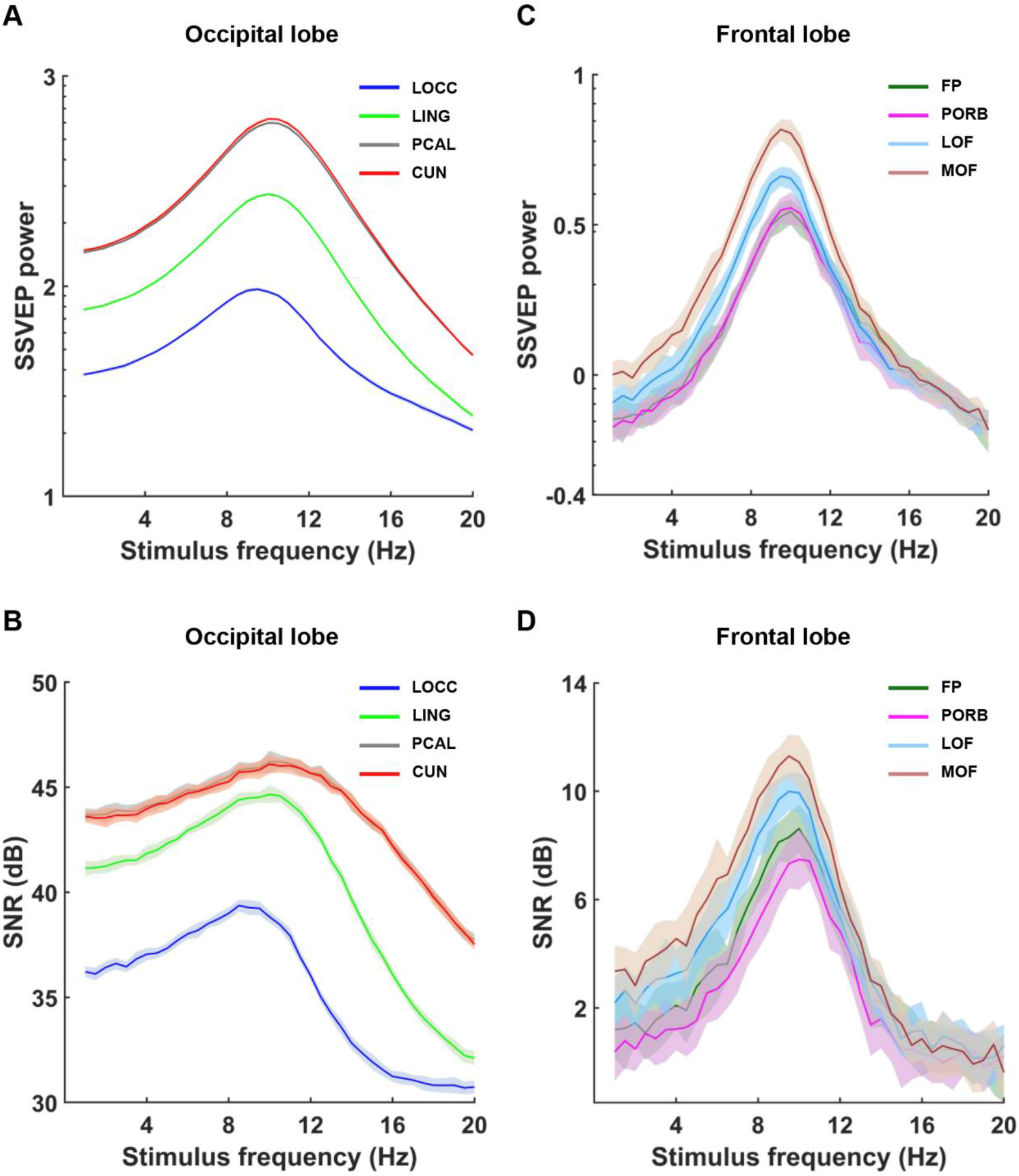
Dependence of SSVEP responses on stimulus frequency. **A, B:** SSVEP power (A) and the SNR value (B) for different occipital regions at different stimulus frequencies. **C, D:** SSVEP power (C) and the SNR value (D) for different frontal regions at different stimulus frequencies. All data are plotted as the mean (curve) ± SD (standard deviation; shaded region). A typical frequency-sensitivity range of 8-12 Hz is present in each region. The occipital and frontal regions showed optimal responses to external periodic visual stimuli in the alpha frequency band (8-12 Hz).

In reality, the performance of the SSVEP responses can also be influenced by the amplitude of the external periodic stimulus. As shown in Fig. 5, positive relationships were observed between SSVEP responses (i.e., SSVEP power and the SNR value) and stimulus amplitude in both the occipital and frontal regions. When the external stimulus was weak, the model generated spontaneous neural activity that was comparable to real electrophysiological recordings. Under this condition, only weak SSVEP responses were detected in the occipital regions (Figs. 5A and 5B). For a large stimulus amplitude, the dynamics of these occipital regions responded well to the external periodic stimulus, thus inducing relatively stronger SSVEP responses (Figs. 5A and 5B). Due to strong interactions among brain regions, evoked neural activity in these occipital regions could be transmitted to the frontal lobe through nerve fibers in a reasonably strong manner. Consequently, large values of both SSVEP power and SNRs were observed in the frontal regions (Figs. 5C and 5D).

**Figure 5.**
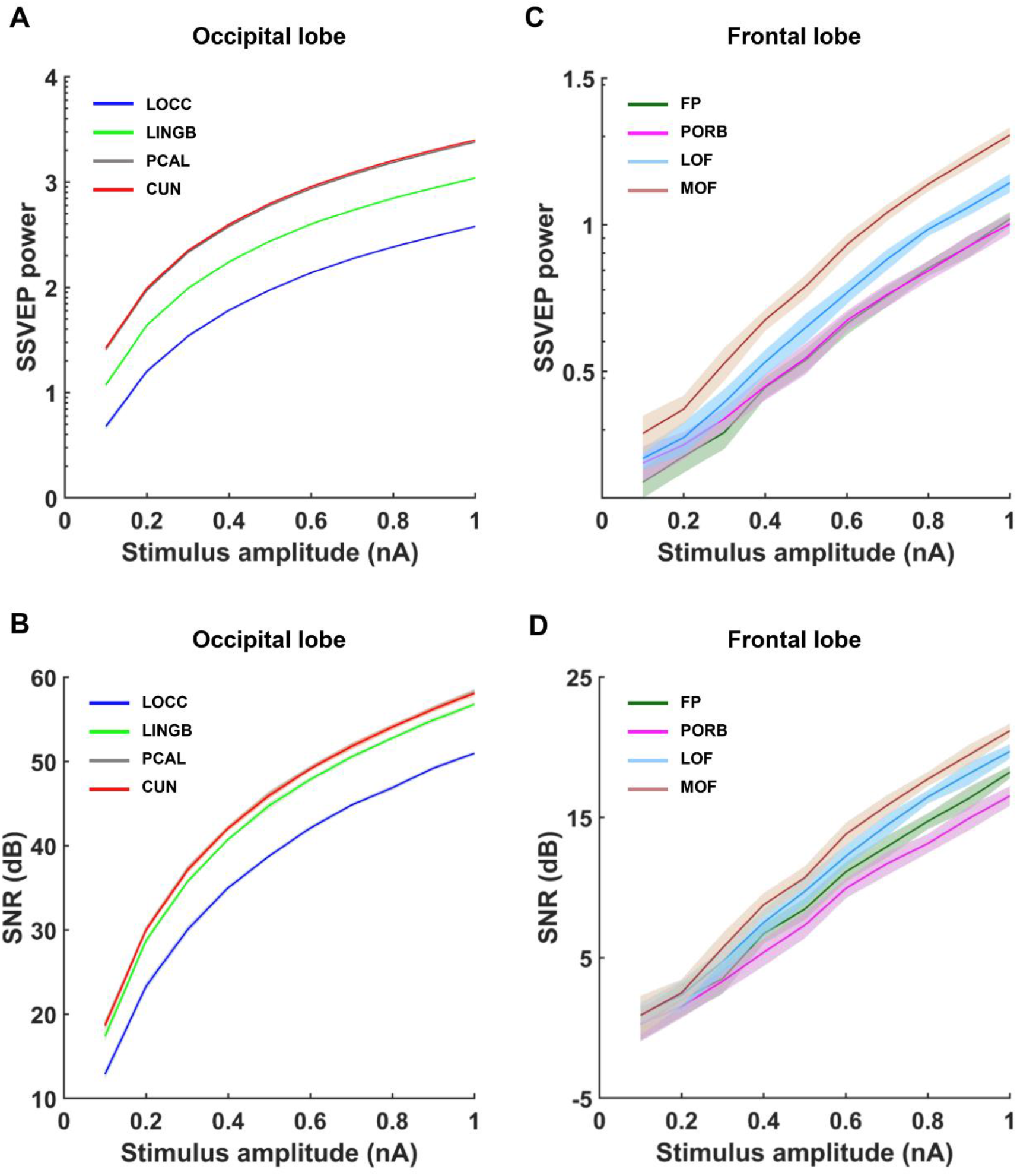
The performance of SSVEP responses is influenced by the stimulus amplitude. **A, B:** SSVEP power (A) and the SNR value (B) for different occipital regions at different stimulus amplitudes. **C, D:** SSVEP power (C) and the SNR value (D) for different frontal regions at different stimulus amplitudes. All data are plotted as the mean (curve) ± SD (shaded region). The positive relationships between SSVEP responses and stimulus amplitude can be observed in both the occipital and frontal regions.

### 3.4 Dynamical nature of the frequency sensitivity of SSVEPs

Exploring the dynamic nature of frequency-sensitivity behavior can deepen our mechanistic understanding of SSVEPs. Intuitively, we hypothesized that the frequency sensitivity of SSVEPs may be caused by both the entrainment and resonance due to the cooperation of intrinsic brain oscillations and external periodic stimuli. In physics, entrainment reflects that the natural oscillation of an internal oscillator perturbed by an external periodic stimulus becomes synchronized to the periodic driven force, whereas resonance describes the phenomenon of increased amplitude that occurs when the frequency of a periodically applied stimulus is equal or close to an intrinsic frequency of the system on which it acts. To examine whether our hypothesis is true, we simulated large-scale brain dynamics in the resting state and stimulus-evoked state. In Fig. 6A, we compared the average power spectral density of the whole brain between the resting state (black dotted line) and stimulus-evoked state (colored lines). At the resting state, the brain dynamics generated by the model showed relatively strong powers in the alpha band, which matched well with the frequency-sensitivity range (8-12 Hz) of SSVEPs. When the model is driven by an external periodic stimulus, the peak frequency of neural oscillations is shifted to the stimulus frequency (colored lines). Further analysis showed that the collective neural activity among different brain regions exhibited relatively strong synchronization when the stimulus frequency was in the alpha band (Fig. 6B). This evidence indicates the occurrence of neuronal entrainment and such entrained oscillation tends to be weakened provided that the stimulus frequency and intrinsic oscillation frequency are mismatched. Moreover, we also observed the increased amplitude of neural oscillations in the occipital lobe when the stimulus frequency of the external periodic input is close to the intrinsic oscillation frequency of the resting-state brain dynamics (Supplementary Fig. 2). This implies that resonance may also contribute to the origin of SSVEPs and, under such condition, the strongest SSVEP power can be detected at the whole-brain level (yellow line in Fig. 6A). To a certain extent, such resonance-induced enhancement in SSVEP response might also impact the frequency sensitivity of SSVEPs. Overall, these findings supported our hypothesis that the frequency sensitivity of SSVEPs might be determined by the combined effects of neuronal entrainment and resonance.

**Figure 6.**
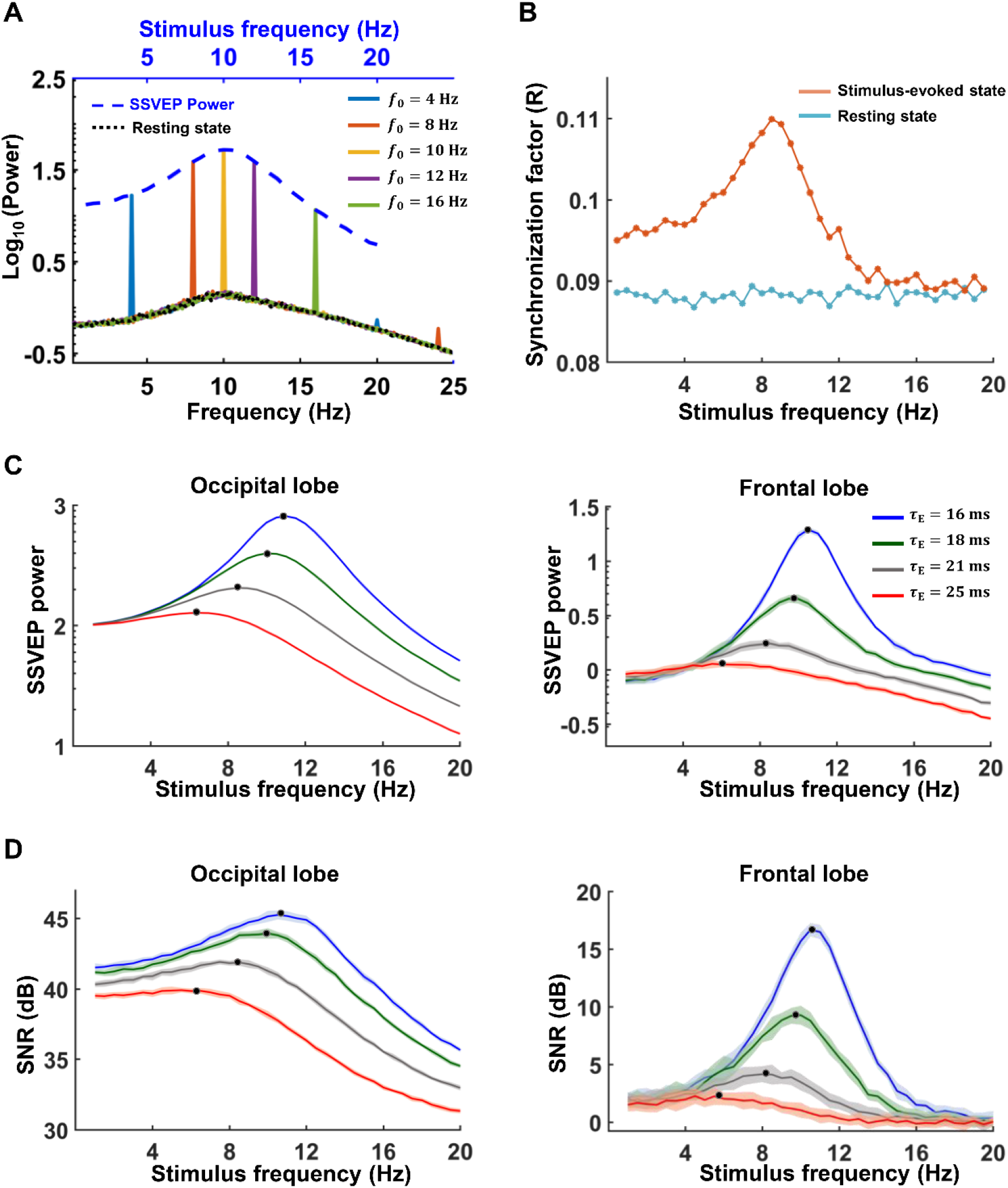
Dynamical nature of the frequency sensitivity of SSVEPs. **A:** The average power spectrum density across all brain regions in the resting state (black dotted line) and in different stimulus-evoked states (colored lines). The blue, orange, yellow, purple, and green lines represent the average power spectrum at stimulus frequencies of 4 Hz, 8 Hz, 10 Hz, 14 Hz, and 16 Hz, respectively. For comparison, we also plotted the SSVEP power (blue dotted line) as a function of stimulus frequency in the same figure. **B:** The synchronization factor *R* for brain networks at both the resting and stimulus-evoked states under different frequencies. **C:** The average SSVEP power (C) and the average SNR (D) for the occipital (left) and frontal (right) lobes at different stimulus frequencies. Here, different colors in C and D represent different time constants of excitatory neural populations, and black dots indicate curve peaks.

A naturally arising question is whether the frequency-sensitivity range of SSVEPs can be modulated by endogenous factors in the brain. We argue that this is possible because such a frequency-sensitivity range should be changed with the intrinsic oscillation frequency of the brain. To test this notion, we illustratively varied the intrinsic oscillation frequency of the large-scale brain model by tuning the default time constant of excitatory neural populations *τ*_*E*_ (Figs. 6C and 6D). This parameter determines how quickly the firing rate of an excitatory neural population decays to the baseline level of spontaneous brain activity. Theoretically, the increase in the time constant of excitatory neural populations slowed down the model dynamics and thus resulted in a low intrinsic oscillation frequency. For both occipital and frontal lobes, such a decreasing effect on intrinsic oscillation frequency shifted the frequency-sensitivity range of SSVEPs toward the left, corresponding to a stimulus within a low-frequency regime (Figs. 6C and 6D). By reducing the time constant of excitatory neural populations, the opposite results were obtained because of the emergence of a high intrinsic oscillation frequency (Figs. 6C and 6D). Indeed, a similar frequency-sensitivity modulation of SSVEPs could also be observed by tuning the time constant of inhibitory neural populations (data not shown) or other endogenous factors that impact the intrinsic oscillation frequency. These results provide evidence that the frequency-sensitivity range of SSVEPs may change together with the intrinsic oscillation frequency of the brain.

### 3.5 Network properties contribute to the performance of SSVEP responses

Given that SSVEPs are regulated by multiple brain areas, we performed graph analysis for brain networks under both resting and stimulus-evoked states. Figs. 7A and 7B show the clustering coefficient and characteristic path length of the evoked brain networks, respectively, at each stimulus frequency (red lines). For comparison, we also plotted these two network measurements for the resting-state brain networks in Figs. 7A and 7B (blue lines). A bell-shaped curve was observed for the clustering coefficient (Fig. 7A), whereas the characteristic path length exhibited an inverted bell-shaped curve (Fig. 7B). Slightly different from SSVEP responses, the large clustering coefficients and small characteristic path lengths mainly appeared in the low alpha band (8-10 Hz). At all frequency points, we found that the stimulus-evoked brain networks displayed stronger clustering coefficients and smaller characteristic path lengths than those of the resting-state brain networks. According to complex network theory, this implied that the evoked brain state may be endowed with a higher small-worldness, thus exhibiting a more efficient FC with high parallel information transfer at the neural activity level. Further examination showed that both global and local efficiency exhibited bell-shaped curves; brain networks evoked by stimuli also presented higher global and local efficiency than the global and local efficiency observed in the resting state (Figs. 7C and 7D). For both types of brain networks, global and local efficiency also achieved their optimal performance when the stimulus frequency was near 9.5 Hz (Figs. 7C and 7D), further supporting that the frequency sensitivity of SSVEPs is determined by the intrinsic oscillation frequency of the brain at the network level.

**Figure 7.**
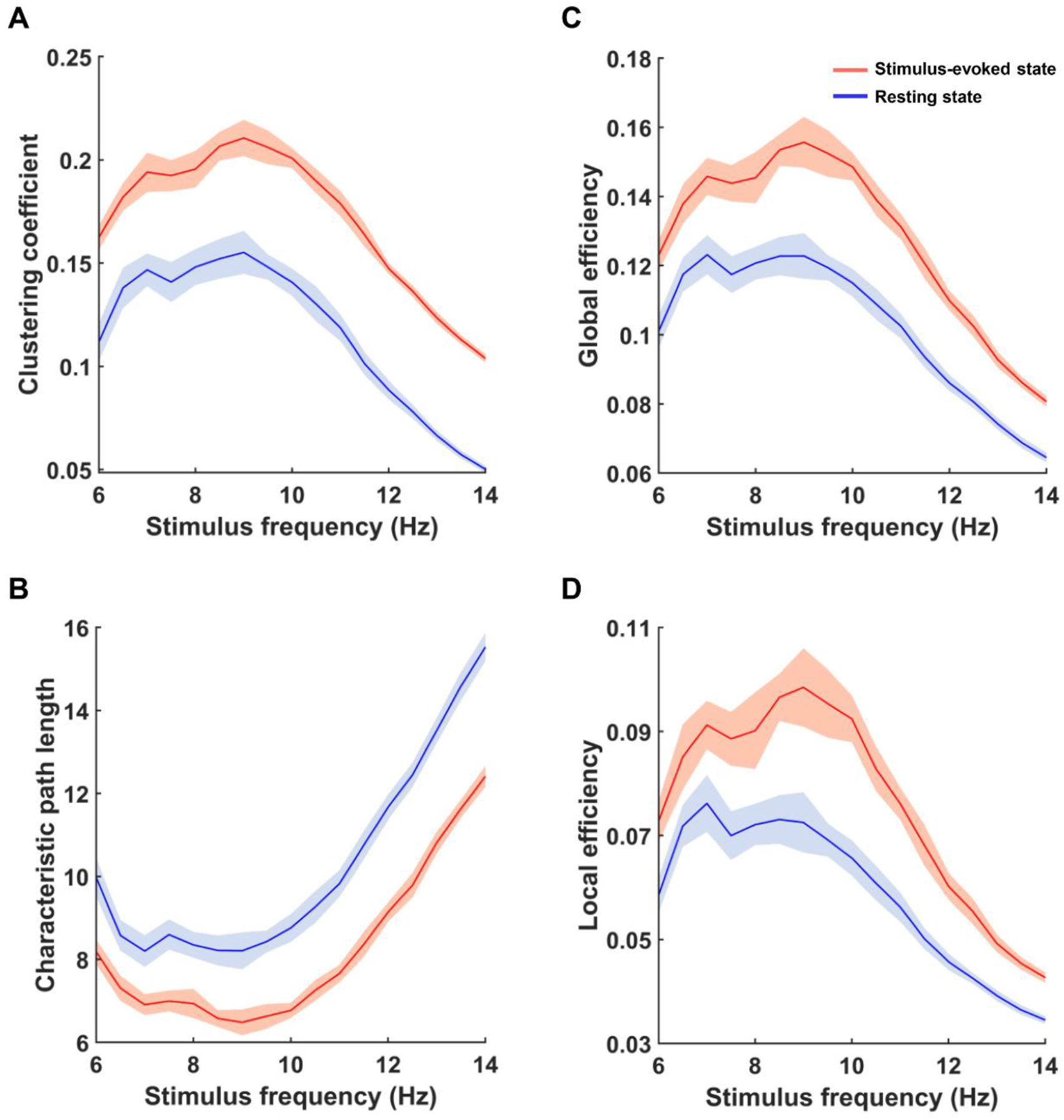
Network properties of the stimulus-evoked and resting brain states at different frequency points. **A-D**: Clustering coefficient (A), characteristic path length (B), global efficiency (C), and local efficiency (D). Red lines indicate the stimulus-evoked network properties at the stimulus frequency point, and blue lines represent resting-state network properties at the corresponding frequency point. All data are plotted as the mean (curve) ± SD (shaded region). The stimulus-evoked brain networks displayed stronger clustering coefficients, global efficiency, and local efficiency and smaller characteristic path lengths than those of the resting-state brain networks.

To gain a better mechanistic understanding of SSVEPs, we also assessed the differences in FC between the stimulus-evoked state and the resting state at the neural activity level by a two-sample Student’s t-test with a significance level of *p* < 0.05 (familywise error (FWE) correction). Compared with resting-state brain networks, no significant decreases in connections were found in networks in the stimulus-evoked state (Fig. 8A). For both low and high stimulus frequencies, enhanced connections in the stimulus-evoked state mostly appeared between the occipital and temporal lobes (see 6 Hz and 14 Hz; Fig. 8A). Under this condition, the evoked neural activity could not be well transmitted to other brain lobes, thus causing weak SSVEP responses in the frontal lobe (Figs. 4C and 4D). For an appropriate stimulus frequency of 9.5 Hz, we identified a broad enhancement in connectivity within the whole brain (Fig. 8A). This resulted in highly efficient FC at the neural activity level corresponding to better SSVEP performance. There might be two possible contributors to this observation: increased neural activity and enhanced network synchronization. By comparing activation between these two brain states, we found that the average firing rates of most brain regions were not changed dramatically in the stimulus-evoked brain state (Fig. 8B). Significantly increased neural activity was only observed for occipital regions because of the direct provocation by the external periodic stimulus (Fig. 8B). Our observation thus provided evidence to rule out the first contributor. Accordingly, such high-efficiency FC was supposed to be a result of the enhanced synchronous neural activity among brain regions (Fig. 6B and Figs. 7A-7D).

**Figure 8.**
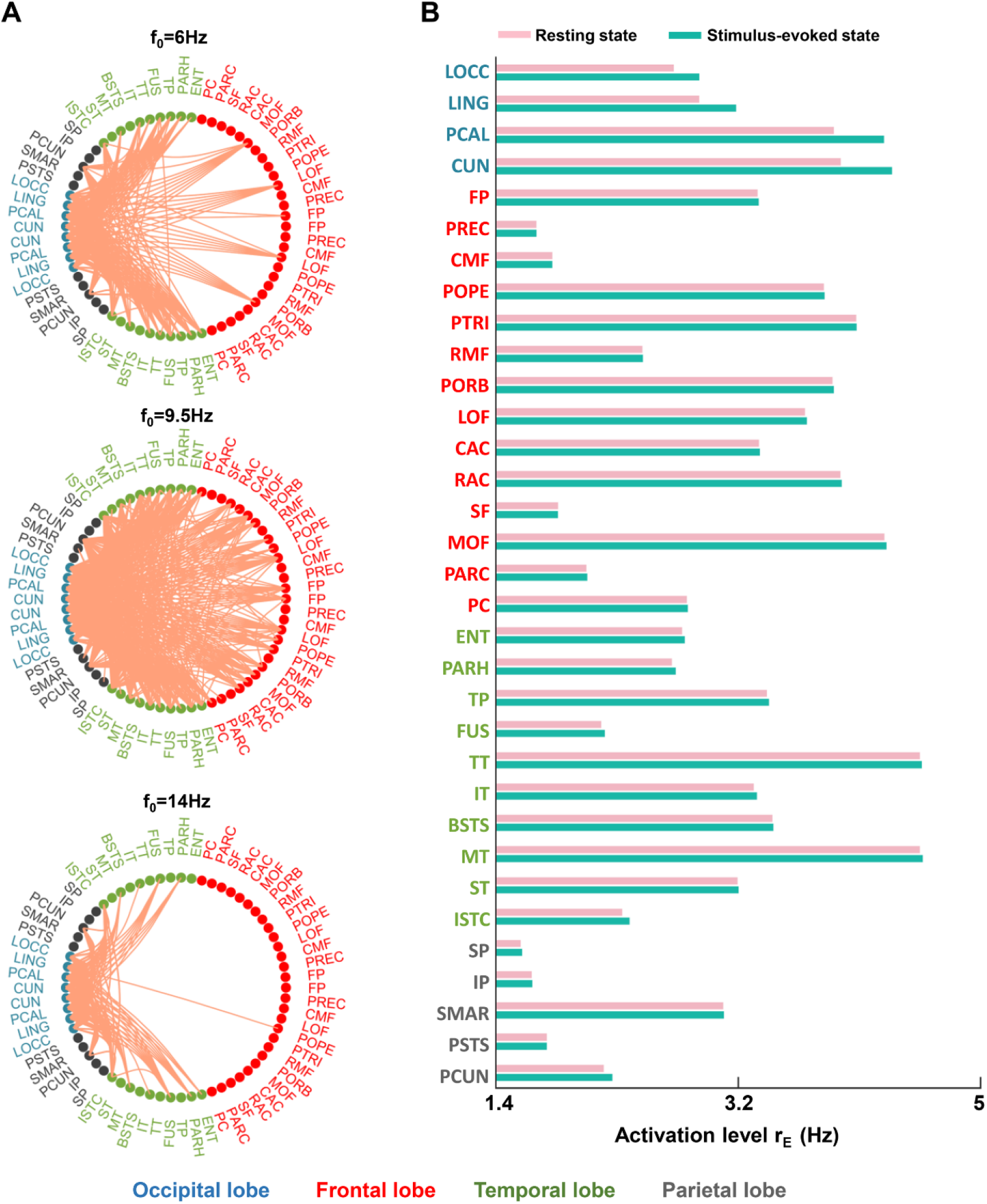
Alterations in connectivity and activation level between the stimulus-evoked and resting brain states. **A**: Significant changes in FC at the neural activity level for different stimulus frequencies (from top to bottom: *f*_0_ = 6 Hz, 9.5 Hz, and 14 Hz). The results were statistically compared by a two-sample student’s t-test with a significance level of *p* < 0.05 (familywise error (FWE) correction). Orange lines indicate enhanced connections at the stimulus-evoked state, and no significantly decreased connectivity was identified after FWE correction. **B**: Comparison of the average activation level for each brain region between the resting state and stimulus-evoked state (*f*_0_ = 9.5 Hz). Under the appropriate stimulus frequency of 9.5 Hz, the connectivity within the whole brain was enhanced and the average activation levels only for regions in the occipital lobe are observed.

Indeed, our real brain cannot always work under its optimized condition corresponding to optimal global-scale coupling (i.e., *G* = 3.01 for the tolerance level of *T* = 0.125), but it is highly possible to operate near this optimal point due to several factors, such as neural plasticity and individual variability. Obviously, different global-scale couplings will lead to distinct SSVEP responses and network properties. To explore the relationships between SSVEP responses and network properties, we changed the value of *G* around this optimal point in our large-scale brain model, and calculated the average SSVEP responses across all regions and different network properties for each fixed *G*. In Figs. 9A-9D, we summarized the dependence of network properties on the average SSVEP responses. The clustering coefficient, global efficiency, and local efficiency showed significant positive correlations with both SSVEP power and SNR values (Figs. 9A, 9C, and 9D; two-tailed Student’s t-test, *p* < 0.01). In contrast, SSVEP responses were negatively correlated with the characteristic path length of brain networks (Fig. 9B). These data indicated that stronger SSVEP responses of the brain must be supported by more efficient FC that is composed of locally and nonlocally distributed brain regions.

**Figure 9.**
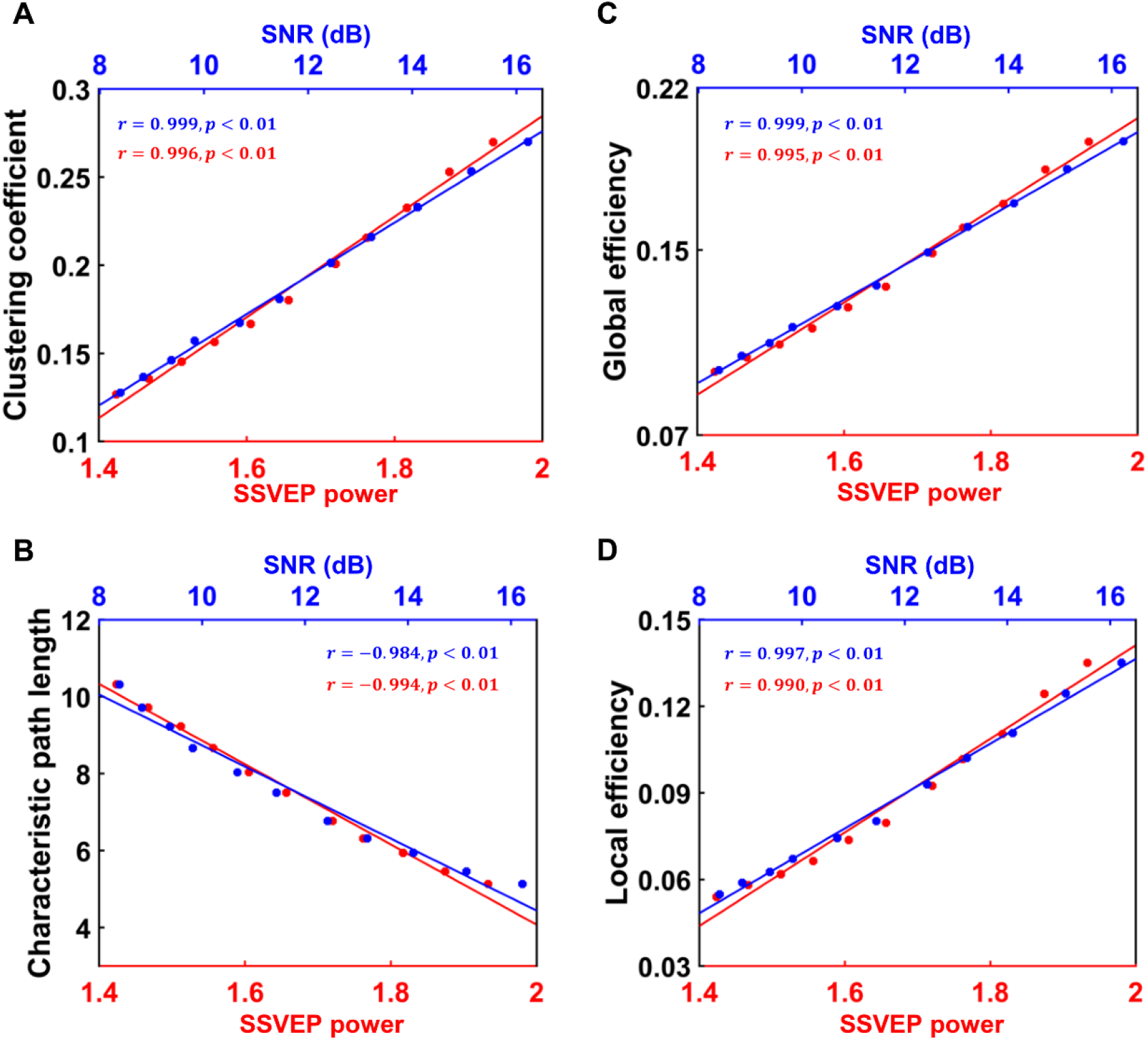
The correlation between SSVEP responses and stimulus-evoked network properties for global-scale couplings near its optimal point of *G* = 3.01. In simulations, the global-scale coupling *G* gradually changed from 2.7 to 3.2, with a fixed step of 0.05. **A-D**: Clustering coefficient (A), characteristic path length (B), global efficiency (C), and local efficiency (D). Blue and red lines represent the SNR value and SSVEP power, respectively. Here, *r* denotes the correlation coefficient, and *p* means the significance level of the correlation coefficient. Statistical significance was determined by the two-tailed Student’s t-test. The clustering coefficient, global efficiency, and local efficiency showed significantly positive correlations with both SSVEP power and SNR values, whereas the SSVEP responses were negatively correlated with the characteristic path length of brain networks.

## 4. Discussion

SSVEPs have been widely used in both neural engineering and cognitive neuroscience, but their underlying dynamic mechanisms within the brain remain to be elucidated. By using a large-scale brain model that integrated multimodal imaging data, we provided computational insights into the mechanistic understanding of SSVEPs at the whole-brain level. Through simulations, we showed that our model can capture the fundamental features of SSVEPs. Under suitable conditions, notable SSVEP responses were detected in both the occipital and frontal lobes, and the performance of SSVEPs was largely impacted by the physical properties of periodic visual stimuli. In particular, we observed that SSVEPs responded optimally to an external periodic stimulus at a specific frequency-sensitivity range of 8-12 Hz (alpha band). Further detailed graph analysis not only revealed that the stimulus-evoked brain network displayed relatively high levels of efficiency and synchronization in a similar frequency-sensitivity range but also confirmed that efficient FC at the neural activity level supports stronger SSVEP responses. Together, these findings contribute to a better understanding of nonlinear SSVEP dynamics in the brain.

The dynamic response of our brain to external periodic input is fundamental for neural information processing (Burkitt et al., 2000; Vialatte et al., 2010). Here, our modeling results indicated that the response of SSVEPs showed the best performance for flickering visual stimulation within 8-12 Hz. Notably, this is in good agreement with previous experimental studies, showing that the largest SSVEP response is elicited by low-frequency visual stimulation in the alpha band (Herrmann et al., 2016; Keitel et al., 2014; Spaak et al., 2014). Our theoretical analysis further revealed that the dynamic nature of the frequency sensitivity of SSVEPs can be attributed to combined effects of nonlinear entrainment and resonance, and the strongest SSVEP response occurs when the stimulus frequency is near the intrinsic oscillation frequency of the brain. As a prominent rhythm of the brain in a resting state, neural oscillations at the alpha band (∼10 Hz) are known to be involved in many types of perceptual or cognitive functions (Herrmann et al., 2016; Pfurtscheller, 2003; Spaak et al., 2014). In particular, alpha-band neural oscillations have been widely detected in EEG recordings across a variety of brain regions, especially during wakeful relaxation with closed eyes (Birca et al., 2006). This suggests that the alpha-dominated rhythm may be the intrinsic neural oscillations of our brain, thus offering a physiological basis in support of the frequency-sensitivity phenomenon observed in SSVEPs.

However, it should be noted that, although several previous modeling studies have also implicated that both entrainment and resonance might serve as a possible mechanism in shaping the frequency response of SSVEPs, most of these studies only used simplified models to simulate the dynamics of local neural populations (Herrmann et al., 2016; Labecki et al., 2016; Notbohm et al., 2016; Roberts & Robinson, 2012). By building a large-scale brain model under constraints of realistic human data, we provided here the first computational evidence that such frequency sensitivity induced by entrainment and resonance can also appear at the whole-brain level.

Our model further predicts that the frequency-sensitivity range of SSVEPs can be regulated by changing the intrinsic oscillation frequency of the brain. At the microscopic scale, a real brain may provide some endogenous mechanisms to automatically adjust intrinsic neural oscillations (Buzsáki & Draguhn, 2004; Herrmann et al., 2016; Latorre et al., 2019); two of these mechanisms are discussed as follows. First, one possibility with high plausibility is directly modulating the intrinsic response properties of neurons. For instance, the decrease in the time constant of cortical neurons in response to decreasing membrane resistance and capacitance tends to result in fast intrinsic neural oscillations (Brunel & Wang, 2003). Under this condition, the frequency-sensitivity range should be shifted toward the high-frequency regime. On the other hand, the concentrations of several types of neurotransmitters have also been found to take part in the regulation of neural oscillations (Basar & Güntekin, 2008; Mariotti et al., 2016). For example, it has been observed that the concentration of gamma-aminobutyric acid (GABA) in the resting state is positively correlated with the frequency of oscillations in response to visual stimulation in humans (Muthukumaraswamy et al., 2009). Therefore, changing the concentrations of several specific neurotransmitters may also provide an alternative approach to modulate the frequency-sensitivity range of SSVEPs. It is worth noting that a similar frequency-sensitivity phenomenon has been extensively reported in neural systems and is believed to play functional roles in highly efficient information processing in the brain (Başar & Güntekin, 2008; Guo et al., 2018). The modulating approaches proposed here may also contribute to a better understanding of these frequency-sensitivity behaviors observed in neural systems.

Highly reliable signal conduction in the brain requires efficient FC. Past experimental studies have revealed that SSVEPs involve both local brain regions and distant, widely distributed brain regions (Birca et al., 2006; Labecki et al., 2016; Zhang et al., 2013). To a certain extent, this might lead to the propagation of SSVEPs in the brain being highly impacted by fundamental properties of cortical networks (Zhang et al., 2013). In the present study, we showed that the performance of SSVEP responses was related to the efficiency of the functional network in the stimulus-evoked state. By comparing network properties at different stimulus frequencies, we identified that the evoked brain state exhibited a relatively highly efficient FC at the neural activity level when the stimulus frequency was in the low alpha band. In this specific stimulus frequency region, we found that more enhanced connections existed between the occipital-temporal and frontal regions compared with the connections noted in other stimulus frequencies, thus ensuring good propagation of SSVEPs in the brain. In addition, our analysis suggested that the emergence of such highly efficient FC was mainly influenced by the enhanced synchronous neural activity among brain regions but not by a significant enhancement in neural activation driven by external periodic stimuli. Using limited specific stimulus frequencies, many experimental studies have also observed that stronger SSVEP responses correspond to more efficient functional networks (Thut et al., 2012; Xu et al., 2013). With the assistance of large-scale brain modeling, we further extended this observation to continuous frequency space. We highlight these findings because they established the linkage between the frequency sensitivity of SSVEPs and the high-level performance of stimulus-evoked brain networks in the low alpha band.

There is a broad consensus that individual differences inevitably exist in many SSVEP studies. In particular, it has been experimentally observed that the responses of SSVEPs display substantial variability across subjects (Koch et al., 2008; Labecki et al., 2016; Zhang et al., 2013). In addition, different subjects may show distinct SSVEP peak frequencies. Notably, our modeling results might provide explainable insights into individual differences observed in experimental studies. On the one hand, the development of the human brain is highly susceptible to changes in a complicated environment (Corbetta et al., 2008; Kramer et al., 2004). During the development of the brain, this factor influences a dynamic change of structure-function relationships for different subjects, thus leading to distinct network efficiency in their FC. As discussed above, the differences in FC efficiency will thus result in substantial variability in SSVEP responses across subjects. On the other hand, alpha-band neural oscillations are believed to contribute the most prominent intrinsic oscillation frequency to the brain (Keitel et al., 2014; Pfurtscheller, 2003; Spaak et al., 2014). In the literature, it has been reported that neural oscillations in the alpha band are highly associated with thalamocortical interactions and that the alpha peak frequency may change with age (Birca et al., 2006; Cantero et al., 2009). Intriguingly, accumulating data have revealed that different subjects may exhibit a certain level of individual variability in alpha peak frequency (Haegens et al., 2014). If our above findings on the frequency sensitivity of SSVEPs can reflect real behavior, such alpha peak variability may provide a physiological basis for the experimentally observed individual differences in SSVEP peak frequency.

In the present study, we only focused on the dynamic mechanisms of SSVEPs and did not involve any cognitive process. Therefore, our model is assumed to be simply driven by the same external stimulus in bilateral occipital lobes. However, a large number of studies have provided evidence that several cognitive processes, such as attention, may take part in the modulation of SSVEP response (Gulbinaite et al., 2019; Hillyard et al., 1997; Keitel et al., 2014; Keitel et al., 2017; Keitel et al., 2019; Müller et al., 1998; Müller & Hillyard, 2000). In particular, both the negative and positive attentional modulation of alpha-band SSVEPs have been widely observed even in the similar experiments (Keitel et al., 2014; Keitel et al., 2017), and such seemingly contradictory findings can be reconciled with different analyzing approaches (Keitel et al., 2019). Several previous studies also indicated that effects of attention on SSVEPs can be observed up to the gamma band, and the sign of attentional modulation of SSVEP amplitude might be frequency dependent (Gulbinaite et al., 2019; Herrmann, 2001). By introducing well-designed stimulus paradigms, our large-scale brain model could be also used to investigate the dynamic mechanisms of attentional modulation of SSVEPs and unify distinct experimental observations in different parameter regimes, another topic that deserves to be explored in the future studies.

Although our large-scale model of the brain is a powerful tool for reproducing the fundamental characteristics of SSVEPs at the system level, we must admit that this model is idealized and can be extended in several aspects. First, we simulated the dynamics for each brain region by using a simplified microcircuit structure composed of a group of excitatory and inhibitory populations. However, the cerebral cortex of the mammalian brain is organized into layers of specialized neuronal subtypes (Burt et al., 2018; Greig et al., 2013; Miller et al., 2019). Previous modeling studies have shown that such laminar specification may perform important functions in signal propagation and modulation between brain regions (D’Souza & Burkhalter, 2017). Therefore, it is reasonable to further construct a more physiological large-scale brain model with a detailed laminar structure and explore how the cortical laminar structure impacts the propagation of this evoked neural activity in the brain. Second, we did not incorporate the transmission delay in our model. Indeed, the transmission delay between two brain regions is highly dependent upon their distance, which may range from several milliseconds to hundreds of milliseconds (Kringelbach et al., 2020; Ziaeemehr et al., 2020). In theory, introducing the distant-dependent time delay into a large-scale brain model will significantly enrich the model dynamics and influence SSVEP responses, a prediction that deserves to be examined in future studies. Finally, we ignored the hierarchical organization of the human brain in the model. By developing a large-scale dynamic model of the macaque neocortex with embedded hierarchy, previous studies have successfully reproduced the functional hierarchy among visual cortical areas that could be compared with experimental observations (Mejias et al., 2016). It has also been proposed that the hierarchical structure may play functional roles in the balance between integration and segregation by mediating neural gain (Shine et al., 2018). In future studies, it will be necessary to further explore whether nonlinear SSVEP dynamics can also be modulated by the hierarchical organization of the brain.

To summarize, we performed a systematic study on the dynamic mechanisms of SSVEPs with a large-scale brain model constrained by empirical human MRI data. We demonstrated that such a biophysical-based model could capture the fundamental features of SSVEP dynamics and reproduce the distributed characteristics of SSVEPs in the brain. Our results indicated that the dynamic nature of SSVEPs is a consequence of neuronal entrainment and resonance, and revealed that the efficient stimulus-evoked FC that emerges in a frequency-sensitivity range near the alpha band contributes to the high-level performance of SSVEP responses. These findings might not only deepen our current understanding of the biophysical mechanisms of SSVEPs but may also inspire testable hypotheses for future experiments. Additionally, our study emphasizes that large-scale brain modeling is a promising approach with a bright future to characterize the dynamics and functions of the brain in continuous parameter spaces under both normal and abnormal states. Further establishing the large-scale brain model at the individual level will form the personalized digital twin brain (DTB), which will greatly promote the applications of virtual brain technology in the studies of individualized medicine.

## Data and Code Availability

All MRI data used in this study can be downloaded from the Github repository of TVB-data (https://github.com/the-virtual-brain/tvb-data/tree/master/tvb_data/connectivity). Codes of the large-scale brain model were developed by Daqing Guo’s Group at the University of Electronic Science and Technology of China, and will be also available by request after the acceptance of this manuscript.

## CRediT authorship contribution statement

Ge Zhang: Methodology, Formal analysis, Visualization, Writing-Original draft preparation. Yan Cui: Formal analysis, Visualization. Yangsong Zhang: Formal analysis, Funding acquisition. Hefei Cao: Methodology Guanyu Zhou: Methodology. Haifeng Shu: Data curation, Methodology. Dezhong Yao: Investigation, Conceptualization, Funding acquisition, Writing-Reviewing & Editing. Yang Xia: Funding acquisition. Ke Chen: Conceptualization, Resources. Daqing Guo: Conceptualization, Supervision, Funding acquisition, Writing-Reviewing & Editing.

## Acknowledgments

We sincerely thank Prof. Peng Xu and Prof. Mingming Chen for the valuable discussions. This work is supported by the National Natural Science Foundation of China (Nos. 31771149, 81861128001, 82072011, 61871423), the Sichuan Science and Technology Program (No. 2018HH0003), and the CAMS Innovation Fund for Medical Sciences (CIFMS) (No. 2019-I2M-5-039).

